# Evaluating the feasibility of medium chain oleochemical synthesis using microbial chain elongation

**DOI:** 10.1101/2024.05.06.592817

**Authors:** Ethan Agena, Ian M. Gois, Connor M. Bowers, Radhakrishnan Mahadevan, Matthew J. Scarborough, Christopher E. Lawson

## Abstract

Chain elongating bacteria are a unique guild of strictly anaerobic bacteria that have garnered interest for sustainable chemical manufacturing from carbon-rich wet and gaseous waste streams. They produce C_6_-C_8_ medium-chain fatty acids which are valuable platform chemicals that can be used directly, or derivatized to service a wide range of chemical industries. However, the application of chain elongating bacteria for synthesizing products beyond C_6_-C_8_ medium-chain fatty acids has not been evaluated. In this study, we assess the feasibility of expanding the product spectrum of chain elongating bacteria to C_9_-C_12_ fatty acids, along with the synthesis of C_6_ fatty alcohols, dicarboxylic acids, diols, and methyl ketones. We propose several metabolic engineering strategies to accomplish these conversions in chain elongating bacteria and utilize constraint-based metabolic modelling to predict pathway stoichiometries, assess thermodynamic feasibility, and estimate ATP and product yields. We also evaluate how producing alternative products impacts the growth rate of chain elongating bacteria via resource allocation modelling, revealing a trade-off between product carbon length and class versus cell growth rate. Together, these results highlight the potential for using chain elongating bacteria as a platform for diverse oleochemical biomanufacturing and offer a starting point for guiding future metabolic engineering efforts aimed at expanding their product range.

**Graphical Abstract:** In this work, the authors use constraint-based metabolic modelling and enzyme cost minimization to assess the feasibility of using metabolic engineering to expand the product spectrum of anaerobic chain elongating bacteria.

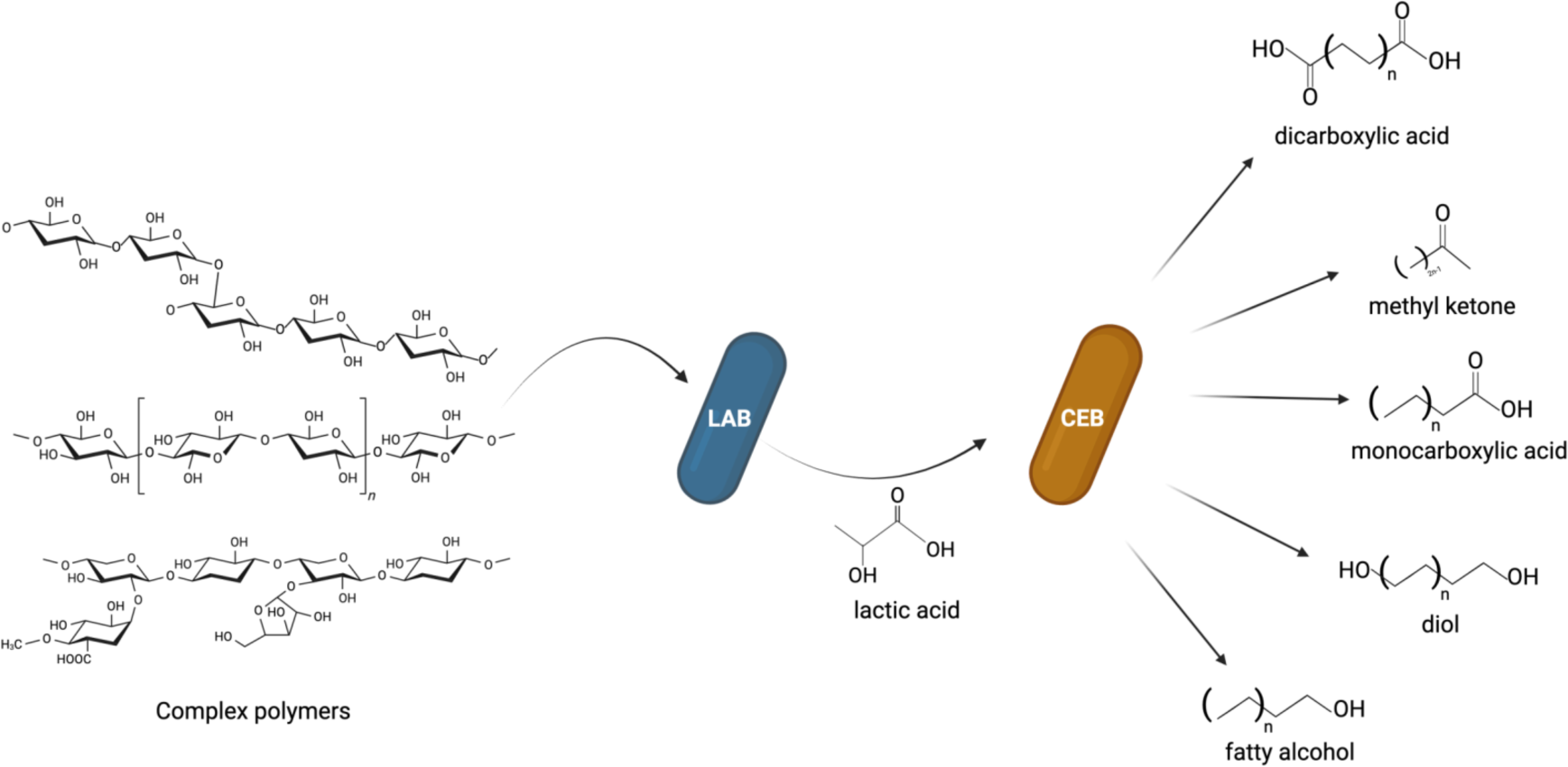

**One Sentence Summary:** In this work, the authors use constraint-based metabolic modelling and enzyme cost minimization to assess the feasibility of using metabolic engineering to expand the product spectrum of anaerobic chain elongating bacteria.

## Introduction

Circular economies require the recycling of societal “wastes” into new bioproducts to support sustainable human activity. Anaerobic fermentation processes can upcycle carbon from agricultural residues, food waste, and industrial off-gases into useful fuels, chemicals, and materials. One promising process is microbial chain elongation, which uses anaerobic microbiomes in open-culture systems to synthesize medium-chain fatty acids (MCFAs) from complex organic and gaseous waste streams (Angenent et al., 2016; Holtzapple et al., 2022; Scarborough et al., 2022). The process works by producing key intermediates, such as lactate and/or ethanol, through either organic waste fermentation (breaking down) or gas fermentation (building up), which subsequently undergo a secondary fermentation to produce MCFAs via chain elongation. Several studies have demonstrated stable MCFA production from organic waste (Grootscholten et al., 2014; Stamatopoulou et al., 2020) and gaseous feedstocks (Bäumler et al., 2022; Diender et al., 2016; Fernández-Blanco et al., 2022) via chain elongation at the bench- and pilot-scale; and more recently a demonstration plant has been built in the Netherlands by the Dutch company, ChainCraft (https://www.chaincraft.nl/).

At the heart of the process is a functional guild of obligate anaerobes called “chain elongating bacteria” that use a native reverse beta-oxidation (RBO) pathway (see Figure 1B) to ferment lactate, ethanol, and other electron-rich organic substrates (e.g., sugars, glycerol) into C_4_-C_8_ carboxylates (i.e., butyrate, hexanoate, and octanoate) as part of their growth. The pathway is unique because it allows redox balancing, while also conserving energy through a novel flavin-based electron bifurcation (FBEB) mechanism (Buckel & Thauer, 2018; Li et al., 2008). Almost all known chain elongators belong to the phylum *Bacillota* (previously *Firmicutes*) and are a phylogenetically and physiologically diverse group. While most chain elongators remain uncultivated, over 15 strains have been isolated and sequenced to date, with most new isolates being reported in the last 10 years (Candry & Ganigué, 2021). These isolates, particularly *Clostridium kluyveri,* have been used to establish synthetic co-cultures to convert gaseous or sugar-based feedstocks to MCFAs and their corresponding alcohols (Bäumler et al., 2022; Diender et al., 2016; Haas et al., 2018; Otten et al., 2022; R. Lynd et al., 2022) which could be expanded to more complex feedstocks by selecting lactate and/or ethanol-producing partners with improved hydrolytic capabilities.

**Figure 1-.**
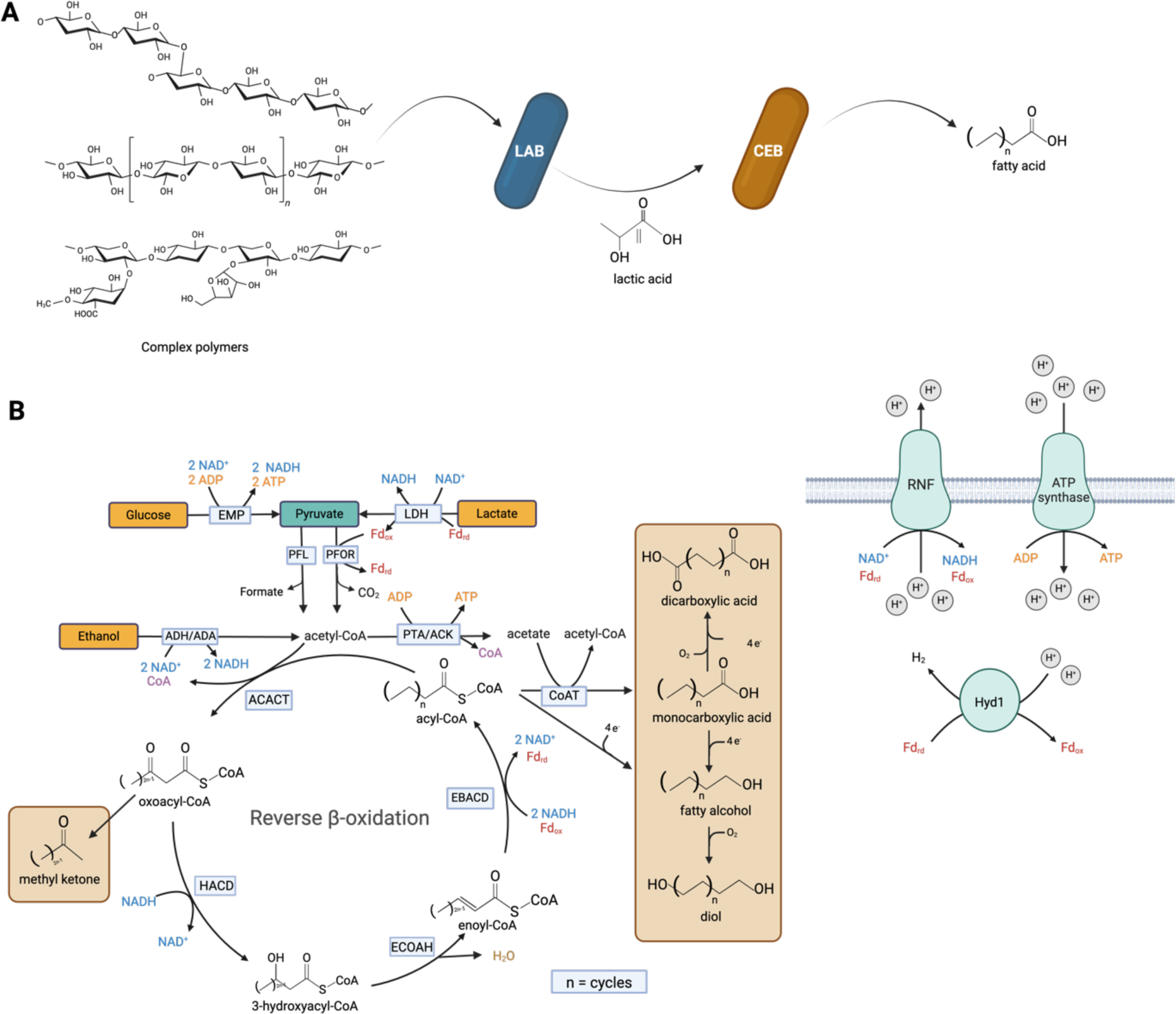
(A) Anaerobic conversion of organic waste to oleochemicals using synthetic co-cultures inspired by natural systems. Complex polymers are degraded by lactic acid bacteria (LAB), which in turn provide lactic acid to chain-elongating bacteria for medium-chain fatty acid synthesis. (B) The reverse beta-oxidation pathway in chain elongating bacteria and proposed product spectrum expansion. EMP: Embden-Meyerhof-Parnas glycolysis; LDH: lactate dehydrogenase; ADH: alcohol dehydrogenase; ADA: acetaldehyde dehydrogenase; PFOR: pyruvate:ferredoxin oxidoreductase; PFL: pyruvate formate lyase; PTA: phosphate acetyltransferase; ACK: acetate kinase; ACACT: acetyl-CoA C acetyltransferase; HACD: 3-hydroxyacyl-CoA dehydrogenase; ECOAH: enoyl-CoA hydratase; EBACD: electron bifurcating acyl-CoA dehydrogenase; CoAT: CoA transferase; Fd: ferredoxin; RNF: proton translocating ferredoxin:NAD^+^ oxidoreductase complex; HYD: Fe-Fe hydrogenase. Created with BioRender.com.

The development of synthetic co-cultures, inspired by mixed culture chain elongation processes, could represent a platform for improving titres, rates, and yields of MCFA production (Figure 1A). Moreover, there is growing interest in genetically modifying the native RBO cycle in chain elongators to expand the product spectrum beyond C_4_-C_8_ monocarboxylates, including alcohols, diols, dicarboxylates, and other bulk chemicals (Agena et al., 2023; Guss & Riley, 2021; Strik et al., 2022). This would enable the anaerobic synthesis of diverse medium-chain oleochemicals from wet and gaseous waste streams, which is expected to have improved economic feasibility compared to sugar-based aerobic fermentations (Holtzapple et al., 2022). However, the technical feasibility of making products other than C_4_-C_8_ monocarboxylates via metabolic engineering of chain elongators remains unexplored. As tools to genetically modify chain elongators emerge (Agena et al., 2023; Cheng et al., 2019; Guss & Riley, 2021), methodologies to intelligently design these new biocatalysts are needed.

In this study, we use constraint-based metabolic modelling along with thermodynamic analyses to evaluate the feasibility of synthesizing diverse medium-chain oleochemicals (C_6_-C_12_ fatty acids, primary alcohols, dicarboxylates, diols, and methyl ketones) from key intermediate substrates (lactate, ethanol, sugars, glycerol) using anaerobic chain elongating bacteria (Figure 1). We first analyze MCFA production scenarios beyond natural C_8_ production up to C_12_, highlighting trade-offs between ATP yield and expected growth rate using a combination of metabolic modelling and enzyme cost minimization analyses. Subsequently, we propose several modifications to the RBO cycle to synthesize target medium chain oleochemical products and use constraint-based metabolic modelling to determine overall pathway stoichiometry, thermodynamic feasibility, and theoretical product yields. Our results indicate that the metabolic engineering of chain elongating bacteria could enable the anaerobic synthesis of diverse medium-chain oleochemicals at industrially relevant yields. Moreover, we identify potential challenges with engineering the RBO pathway in chain elongators and offer potential solutions to overcome them via metabolic engineering. We anticipate these results will provide a useful starting point for engineering microbial chain elongation to serve as a platform for sustainable chemical manufacturing.

## Results and Discussion

### Impact of MCFA Chain Length on Growth Rate and ATP Yield

#### Modelling Chain Elongation beyond Octanoic Acid

Currently, the products of chain elongation are restricted to butyric and hexanoic acids (C_4_-C_6_), with limited synthesis of octanoic acid (C_8_) (Nelson et al., 2017; Zhu et al., 2017). Genetically modifying chain elongating bacteria to improve selectivity and increase MCFA chain length could improve product yields, while also expanding the process to produce MCFAs with larger markets (C_8_-C_12_).

To assess the feasibility of MCFA production beyond C_8_, we predicted overall pathway stoichiometry and redox balance, free energy change, and theoretical ATP and product yields of C_4_-C_12_ fatty acids (FAs) via parsimonious enzyme usage flux balance analysis (pFBA) with ATP yield set as the objective function using a simplified metabolic model describing core chain elongation metabolism (iFermCell193, see Methods) (Table 1). We focused on lactate utilization by chain elongating bacteria because this is their primary substrate in mixed culture processes converting organic wastes (Figure 1) (Contreras-Dávila et al., 2020). Model simulations with different electron donors, including ethanol, glucose, xylose, and glycerol FAs were also evaluated (see Supplemental Material) to demonstrate process feasibility for a range of other possible operating scenarios.

**Table 1.** Predicted stoichiometry for the synthesis of C_4_-C_12_ FAs from lactate.

| Chain Length | Overall Equation | ATP Yield (mol ATP / mol Lactate) | $\Delta G_r^{\circ}$ (kJ/mol) | $\Delta G_r^{\circ}$ per ATP (kJ/mol) | Yield (g FA / g Lactate) |
| --- | --- | --- | --- | --- | --- |
| C<sub>4</sub> | Lactate + 1/2 H<sup>+</sup> → 1/2 Butyrate + CO<sub>2</sub> + H<sub>2</sub> | 0.250 | -28.98 | -160.11 | 0.49 |
| C<sub>6</sub> | Lactate + 2/3 H<sup>+</sup> → 1/3 Hexanoate + CO<sub>2</sub> + 2/3 H<sub>2</sub> + 1/3 H<sub>2</sub>O | 0.333 | -48.53 | -255.34 | 0.43 |
| C<sub>8</sub> | Lactate + 3/4 H<sup>+</sup> → 1/4 Octanoate + CO<sub>2</sub> + 1/2 H<sub>2</sub> + 1/2 H<sub>2</sub>O | 0.375 | -48.53 | -170.22 | 0.40 |
| C<sub>10</sub> | Lactate + 4/5 H<sup>+</sup> → 1/5 Decanoate + CO<sub>2</sub> + 2/5 H<sub>2</sub> + 3/5 H<sub>2</sub>O | 0.400 | -231.56 | -804.43 | 0.38 |
| C<sub>12</sub> | Lactate + 5/6 H<sup>+</sup> → 1/6 Dodecanoate + CO<sub>2</sub> + 1/3 H<sub>2</sub> + 2/3 H<sub>2</sub>O | 0.417 | -195.82 | -483.10 | 0.37 |

The model predicted that ATP yield and overall reaction-free energy increases with MCFA chain length from C_4_-C_12_ (Table 1), indicating MCFA synthesis beyond C_8_ should be theoretically possible. Moreover, the production of C_4_-C_12_ chain lengths from lactate released ≥ 50 kJ/mol indicating these reactions produce sufficient energy for ATP synthesis. C_9_-C_12_ carboxylate production with other electron donors, including ethanol, glucose, xylose, and glycerol, were also found to be feasible (see Supplemental Material). This observation agrees with past modelling results that suggested hexanoic and octanoic acid production should improve ATP yields from lactate compared to butyric acid production (Scarborough et al., n.d.).

The predicted increase in net ATP production is attributed to a greater flux through the proton translocating ferredoxin:NAD^+^ oxidoreductase complex (RNF) for each turn of the RBO cycle, which leads to greater ATP production via the Ion Motive Force (IMF). In the RBO cycle, each successive elongation generates reduced ferredoxin from the electron-bifurcating acyl-CoA dehydrogenase (EBACD). This ferredoxin goes on to drive RNF to recover some NADH and/or is used by a soluble ferredoxin hydrogenase (HYD1) to evolve H_2_ as a terminal electron sink. As confirmed in the predicted flux distributions, elongation to longer chain lengths increases the total NADH demand required to reduce the acyl chains (HACD and EBACD in Figure 1). Thus, with increasing chain length, a greater proportion of the reduced ferredoxin produced by the EBACD is used to regenerate NADH via RNF, rather than driving HYD1. This ultimately leads to increased ATP production through the IMF along with decreased H_2_ evolution for increasing chain length. These results also indicate that past a certain chain length (>C_12_), H_2_ evolution will cease as all electron equivalents will be required solely for chain elongation.

#### Intracellular Thermodynamic Landscape and Resource Allocation

Our initial modelling analysis suggested that the improved ATP yield with longer chain lengths should naturally select for MCFAs beyond C_8_. However, all pure and mixed culture chain elongation studies have only observed C_4_-C_8_ MCFA products. This points to a potential trade-off between growth yield (or ATP yield) and growth rate (or ATP production rate), resulting from optimal resource allocation. In a model based on resource allocation theory (Supplementary Material; inspired by a similar model developed A. I. Flamholz et al., 2024), where catabolic ATP production and anabolic ATP consumption rates are linearly related to the biomass fractions allocated to either function, RBO enzyme cost per unit ATP production flux and chain elongating bacteria growth rate are inversely correlated (Figure 6). This simplified model captures the fact that a higher growth rate is expected to necessitate both a larger pool of anabolic enzyme and ribosomes (Basan, 2018) and a higher ATP production flux, requiring that catabolic machinery produces ATP faster with less enzyme.

To evaluate the enzyme investment per ATP flux required to produce carboxylates of different chain lengths, we performed Enzyme Cost Minimization (ECM) analysis. ECM places a lower bound on a pathway’s enzyme demand per unit flux (g/(mol/h)) by accounting for the pathway’s length and stoichiometry, reaction thermodynamics, and saturation effects following from the optimal set of metabolite concentrations, given appropriate bounds on those concentrations and parametrization of enzyme kinetics (A. Flamholz et al., 2013). This analysis coupled with ATP yield results from our model showed that minimal enzyme cost per unit ATP flux increases with chain length (Figure 2). This suggests that chain elongating bacteria making longer length products invest more of their proteome into high yield catabolism, at the trade-off of having less catalytic capacity for growth.

**Figure 2.**
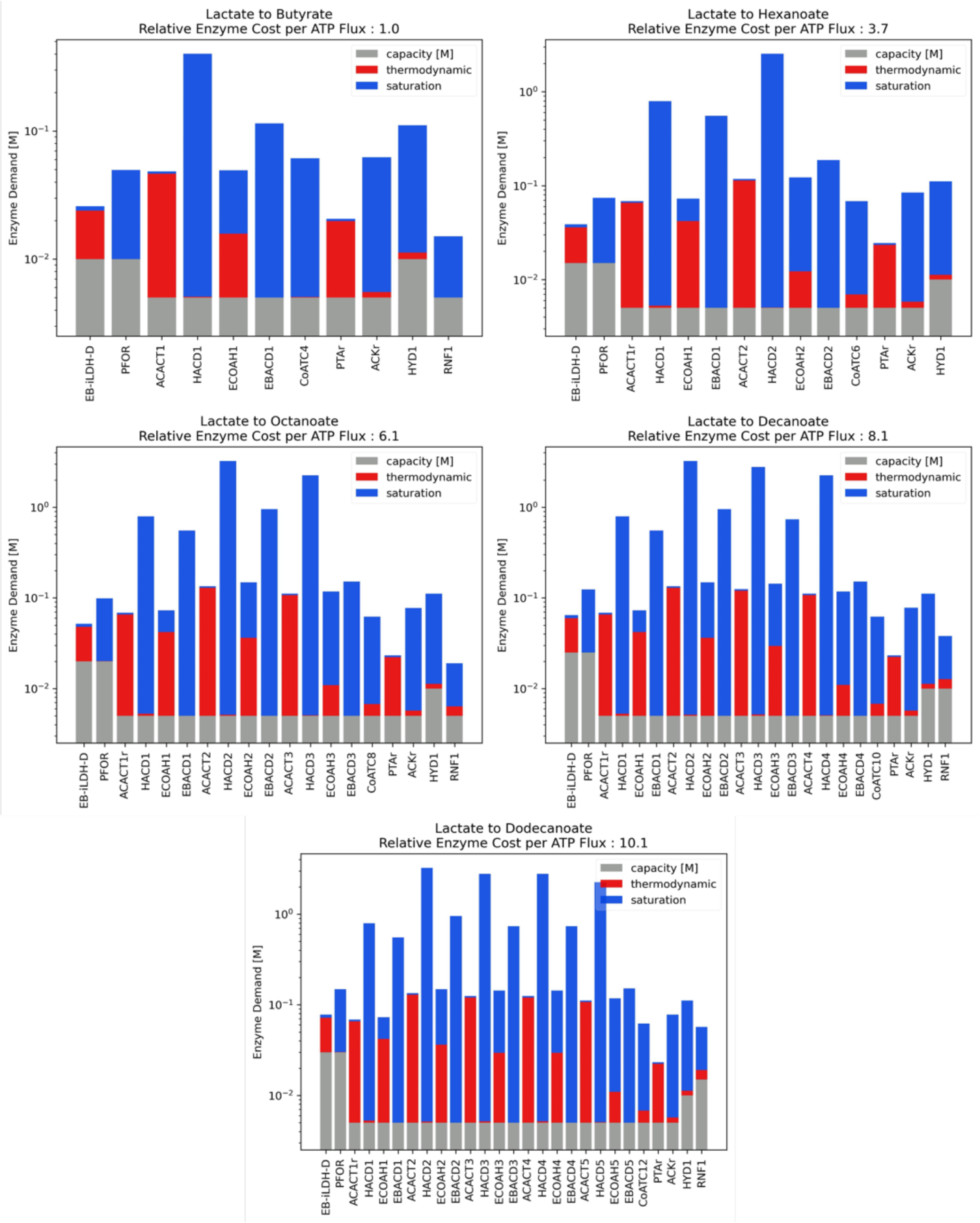
ECM results of RBO for production of carboxylates from lactate. Enzyme cost per unit ATP flux is reported as a multiple of butyrate production’s cost. Each reaction carries a cost which accounts for the pathway’s stoichiometry (grey), increased cost incurred due to a reaction’s proximity to equilibrium (red), and the cost due to sub-saturation reactant concentrations (blue).

According to the optimized free energy changes predicted by ECM, thiolase reactions (ACACT) are the least exergonic in RBO (Supplementary Figure 1). Since these reactions operate near equilibrium, they have a smaller forward to reverse flux ratio (Noor et al., 2014) and therefore require a larger enzyme pool to sustain a given net flux through the pathway. The set of metabolite concentrations which minimizes pathway enzyme cost fixes the thiolase products (oxoacyl-CoAs) at low concentrations to drive these reactions. This heightens the demand for the proceeding enzymes in the pathway, 3-hydroxyacyl-CoA dehydrogenases (HACD) since they are far from saturation (Figure 2). Similarly, enoyl-CoA hydratase reactions (ECOAH) are thermodynamically constrained, meaning their reactant and product concentrations are kept high and low, respectively. This further increases the protein burden of HACD as well as the proceeding electron bifurcating acyl-CoA dehydrogenase (EBACD) (Figure 2). At longer chain lengths, ECOAH reactions are predicted to be more favourable (Supplementary Figure 1), causing the rate at which enzyme cost increases with chain length to decrease.

Interestingly, when oxoacyl-CoA concentrations are constrained by a lower bound of 1 µM, as is the case for all other metabolites, products beyond C_4_ are deemed infeasible due to endergonic subsequent thiolase reactions. To reflect observations of chain elongation beyond C_4_, oxoacyl-CoAs can fall to sub-micromolar levels in this model. An alternative solution, as previously suggested in a similar instance with malonyl dehydrogenase (Noor et al., 2014), is to posit channelling between ACACT and HACD, effectively merging the two into one favourable reaction. Indeed, channelling between ACACT, HACD and ECOAH has been described with crystal structures of bacterial and human beta-oxidation complexes (Ishikawa et al., 2004; Xia et al., 2019). So long as a similar mechanism is active from C_4_ to C_12_, the trend of enzyme cost per ATP flux increasing with chain length is expected to hold. This has major implications for engineering chain elongating bacteria to produce MCFAs beyond C8, as longer chain carboxylates would exhibit higher yields, but at the cost of a lower growth rate. As a result, compensatory efforts to bolster growth rate, for example through adaptive laboratory evolution (Sandberg et al., 2019), may be needed to achieve higher productivity.

### Expanding and Controlling Chain Elongation Products with Metabolic Engineering

Metabolic engineering strategies for the expansion of RBO-derived products have been reviewed for non-chain elongating, model organisms with established genetic tools, such as *Escherichia coli*, *Saccharomyces cerevisiae*, and select acetogenic *Clostridium* species (Tarasava et al., 2022). The approach for these strains relies on the engineered reversal of β-oxidation accomplished through extensive strain engineering such as knockout or repression of native β-oxidation regulators and overexpression of genes from chain elongating bacteria and other oleaginous hosts (Tarasava et al., 2022). Most of these hosts do not natively rely on the RBO cycle for redox balancing or energy conservation which makes the RBO cycle growth-coupled in the chain elongating bacteria. Further, installing orthogonal systems for the engineered reversal of β-oxidation imposes an increased metabolic burden on non-chain elongating hosts as high expression is likely needed to ensure sufficient flux through the pathway (Liu et al., 2018). This can lead to redox limitations or inefficient substrate use for chain elongation in these non-chain elongating hosts and approaches like two-phase production are likely needed to maximize product yields (Lange et al., 2016). Production with non-chain elongating hosts will likely require further strain engineering to improve product yields and selectivity.

As an alternative to engineering model organisms, tools to genetically modify chain elongating bacteria are beginning to emerge (Agena et al., 2023; Cheng et al., 2019; Guss & Riley, 2021), which will open the door to expanded opportunities to produce oleochemicals via anaerobic fermentation. Chain elongating bacteria are ideal hosts for RBO-based bioproduction as the RBO cycle is innately growth-coupled in these strains (no reconstruction needed) and they can utilize solid and gaseous waste feedstocks rather than relying on refined substrates (e.g., sugars). Further, product yields in anaerobes are typically much greater as they do not synthesize as much biomass compared to aerobic systems and most electrons end up in products (Cueto-Rojas et al., 2015). However, the feasibility of pathways for the production of other oleochemicals including, fatty alcohols, dicarboxylates, diols, methyl ketones, has not been explored in anaerobic chain elongating bacteria. Here, we propose potential production pathways and enzymes required to generate a wide range of oleochemicals (Figure 3) and use pFBA and ECM to assess the feasibility and potential effects on ATP yield and growth rate.

**Figure 3.**
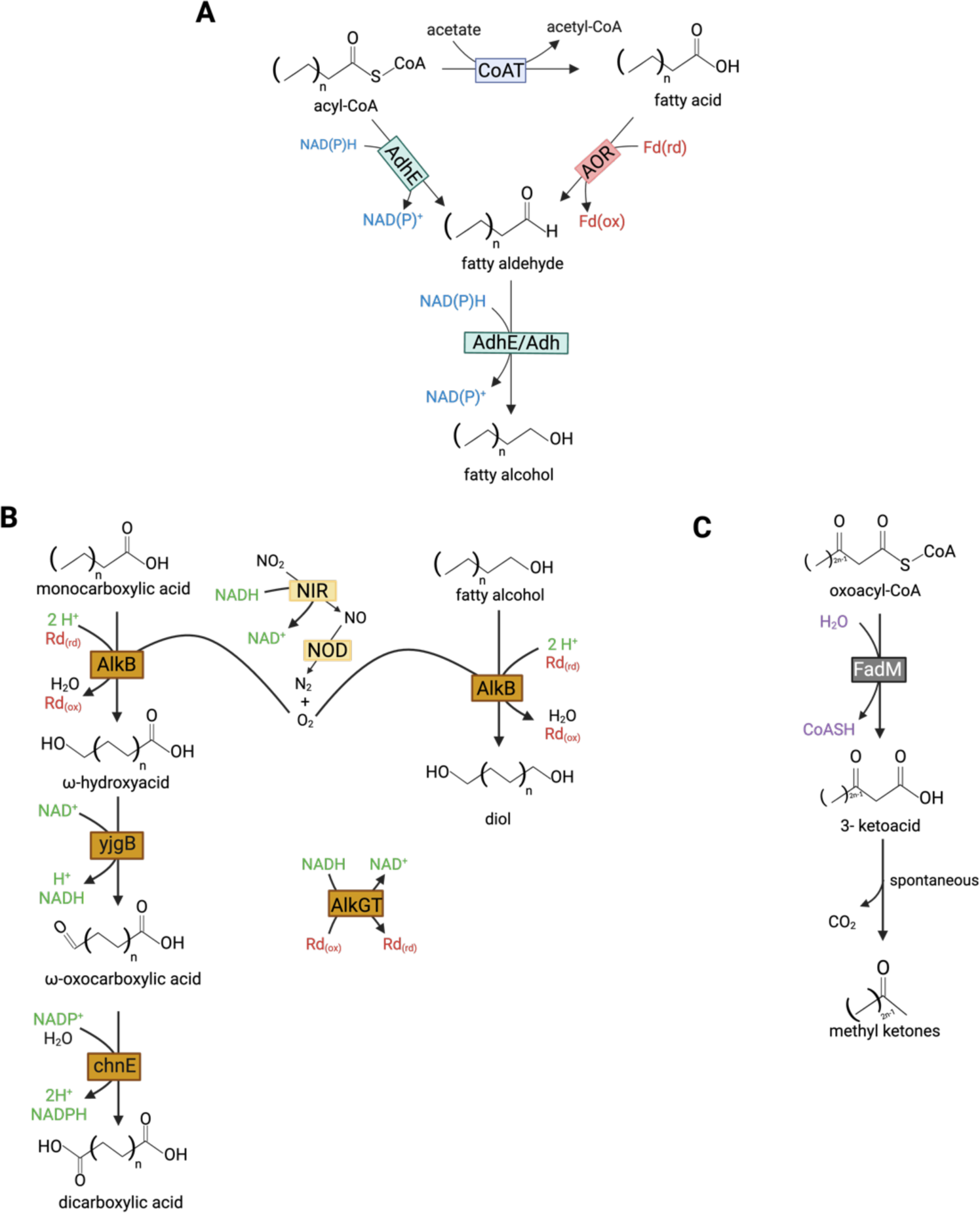
Proposed pathways for chain elongating bacteria product spectrum expansion. A: Fatty alcohols pathway. AdhE: bifunctional aldehyde-alcohol dehydrogenase; AOR: aldehyde:ferredoxin oxidoreductase; FadM: acyl-CoA thioesterase. B. Diol and dicarboxylic acids pathways. NIR: nitrite reductase; NOD: nitric oxide dismutase; AlkBGT: alkane 1-monooxygenase complex; yjgB: cinnamyl alcohol dehydrogenase; chnE: 6-oxohexanoate dehydrogenase; Rd: rubredoxin. C: Methyl ketones pathway. FadM: thioesterase. Created with Biorender.com.

iFermCell193 was further modified to capture the reactions required for hexanoic acid conversion to other products (iFermCell356). Table 2 summarizes the potential pathway stoichiometries for the conversion of hexanoic acid (C_6_) to the corresponding fatty alcohol, dicarboxylate, and diol using lactate as an electron donor, along with the net ATP fluxes and overall Gibbs free energy change of reaction for each substrate-product pair as predicted by our model. All predicted pathway stoichiometries have overall favourable standard Gibbs free energies of reaction and release ≥ 50 kJ/mol per ATP required for ATP production. The model also predicts that net ATP yield is not negatively impacted by conversion into these other bioproducts which indicates that their production can be growth-coupled, similar to MCFAs. In the following sections, we describe potential metabolic engineering strategies to accomplish these conversions in chain elongating bacteria and discuss specific shifts in redox requirements that occur predicted by the model, along with the impact on the metabolic cost of the additional enzymes estimated by ECM.

**Table 2.** Predicted pathway stoichiometry for different C_6_ oleochemicals produced from lactate.

| Product;<br>Pathway | Overall Equation | ATP Yield<br>(mol ATP /<br>mol Lactate) | $\Delta G_r^{\circ}$ | $\Delta G_r^{\circ}$ per<br>ATP | Yield<br>(g P/g S) |
| --- | --- | --- | --- | --- | --- |
| Carboxylate;<br>RBO | $\text{Lactate} + 2/3 \text{ H}^+ \rightarrow 1/3 \text{ Hexanoate} + \text{CO}_2 + 2/3 \text{ H}_2 + 1/3 \text{ H}_2\text{O}$ | 0.333 | -26.75 | -160.11 | 0.43 |
| Alcohol;<br>RBO +<br>AdhE | $\text{Lactate} + \text{H}^+ \rightarrow 1/3 \text{ Hexanol} + \text{CO}_2 + 2/3 \text{ H}_2\text{O}$ | 0.333 | -45.18 | -255.34 | 0.38 |
| Alcohol;<br>RBO + AOR | $\text{Lactate} + \text{H}^+ \rightarrow 1/3 \text{ Hexanol} + \text{CO}_2 + 2/3 \text{ H}_2\text{O}$ | 0.500 | -45.18 | -170.22 | 0.57 |
| Dicarboxylat<br>e; RBO +<br>AlkBGT | $\text{Lactate} + \text{H}^+ + 2/3 \text{ NO}_2^- \rightarrow 1/3 \text{ Adipate} + \text{CO}_2 + 2/3 \text{ H}_2 + \text{H}_2\text{O} + 1/3 \text{ N}_2$ | 0.333 | -228.21 | -804.43 | 0.54 |
| Diol;<br>RBO +<br>AlkBGT | $\text{Lactate} + 5/4 \text{ H}^+ + 1/2 \text{ NO}_2^- \rightarrow 1/4 \text{ 1,6-Hexanediol} + 1/4 \text{ Acetate} + \text{CO}_2 + \text{H}_2\text{O} + 1/4 \text{ N}_2$ | 0.5 | -191.63 | -483.10 | 0.66 |

#### Fatty Alcohol Production

Medium-chain fatty alcohols (FAOHs) could be produced using MCFAs as building blocks. Even though FAOH production pathways (Figure 3A) exist in anaerobic bacteria, such as acetogens and butanol-producing clostridia (Moon et al., 2016), this has yet to be observed in most chain elongating bacteria. In nature, acetogenic bacteria use aldehyde:ferredoxin oxidoreductase (AOR) to convert FAs into fatty aldehydes in monoculture or when they are co-cultured with chain elongating bacteria (Benito-Vaquerizo et al., 2020; Diender et al., 2016). In another instance, aldehyde-alcohol dehydrogenase (AdhE) can convert acyl-CoAs into aldehydes (Mehrer et al., 2018). In these two scenarios, specific electron carriers (1 reduced ferredoxin and 1 NAD(P)H in the former and 2 NAD(P)H in the latter), are necessary to obtain the final product. Final conversion relies on the alcohol dehydrogenase (Adh) activity or the bifunctionality of AdhE (Liew et al., 2017). Both AOR- and AdhE-based FAOH production would result in redox balance shifts if implemented in the chain elongating bacteria as a net redox demand is needed to conduct the reductions.

We incorporated both AOR and AdhE FAOH synthesis pathways in our model and assessed their effects on FAOH production, redox balances, and ATP yield. In chain elongation to FAs, H_2_ is evolved as an electron sink to maintain redox balance. We hypothesized that instead of producing H_2_, these electron equivalents could be used to reduce the carboxylate to the alcohol. Consistent with this, the model predicted that hexanol (C_6_ FAOH) production does not result in H_2_ production (Table 2), and instead, flux is redirected from HYD1 to RNF to produce NADH required by the and AOR and AdhE pathways (Supplementary Material). However, in the AOR pathway, the model predicted a net increase in ATP flux yield through the combined action of IMF and substrate-level phosphorylation as compared to hexanoic acid production (Supplementary Material).

ECM results show that hexanol production via AOR carries an enzyme cost per ATP flux of 2.9 times that of butyrate, whereas hexanoic acid production requires 3.7 times more than butyrate (Figure 4). This reflects the increased ATP yield from FAOH production with AOR and the fact that the RBO cycle reactions are more thermodynamically hindered than redox reactions for FAOH formation from FAs; the additional protein needed to carry flux through the latter represents a small fraction of the pathway’s protein demand. In the AdhE case, ECM results show that hexanoic acid and hexanol production require approximately equal enzyme investments of 3.7 and 3.5 times the cost of butyrate production, due to bypassing the use of PTA and ACK for alcohol production. In both AOR and AdhE cases, it can be expected that chain elongating bacteria engineered for FAOH production will not experience significant growth rate penalties compared to production of MCFA of the same chain length, and in fact, may grow faster if the AOR pathway is implemented.

**Figure 4.**
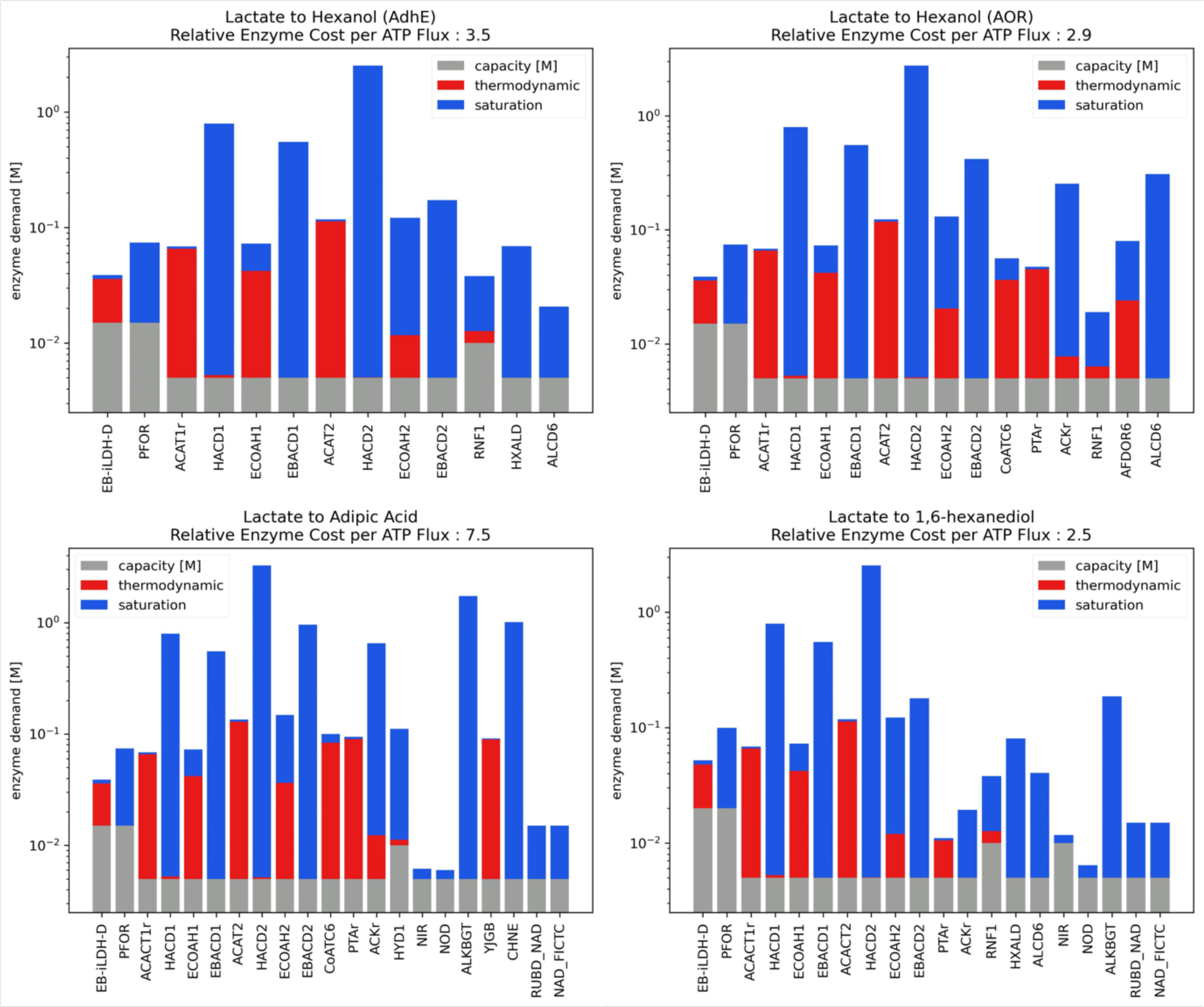
ECM results of RBO for production of novel chain elongation products from lactate. Enzyme cost per unit ATP flux is reported as a multiple of butyrate production’s cost.

#### Dicarboxylate and Diol Production: Pathways Requiring Oxygen

The anaerobic oxidation of MCFAs into diols and dicarboxylic acids poses a significant metabolic engineering challenge in chain elongators, as the lack of oxygen as an electron acceptor results in a considerable decrease in reaction energetics. Methane oxidation is a clear example, where the anaerobic pathway needs to be coupled to nitrite or sulphate reduction (Knittel & Boetius, 2009). Alkane-degrading microbes can anaerobically oxidize C-H bonds by fixing CO_2_ and converting alkanes into FAs. It is hypothesized that this mechanism is initiated by an ethylbenzene dehydrogenase and the whole pathway happens at the great expense of 6 ATP equivalents, suggesting that an unexplored energy-coupling mechanism may take place in these organisms given the low ATP yields in anaerobes (Supplementary Figure 2) (Heider et al., 2016; Shou et al., 2021). Alternatively, alkane monooxygenase complexes like AlkBGT have been used in the hydroxylation of terminal ω-carbons of different compounds, including FAs (Clomburg et al., 2015). However, implementing AlkBGT in anaerobes is currently infeasible due to its requirement for O_2_.

To solve these problems, we envision making use of another hypothesized hydroxylation mechanism in anaerobes: the intra-aerobic pathway (Ettwig et al., 2010). Achieving such conditions requires “oxygen donors”, such as nitrite, hydrogen peroxide, or perchlorate, whose decomposition provides the required oxygen for AlkBGT. Since oxygen presence can be problematic in this scenario, it is important to maintain its intracellular concentrations low, which could be achieved by tuning NIR/NOD and AlkBGT expression or even improving AlkBGT rates through protein engineering. If oxygen levels are still high enough to disrupt cell function, a metabolic strategy that could be explored would be to confine the oxygen generation reaction within microcompartments, pseudo-organelles common in various organisms, including anaerobes, which have been used in several applications, such as protecting cells from toxic intermediates and oxygen (*Bacterial Microcompartments | SpringerLink*, n.d.; Kennedy et al., 2021).

In our proposed pathway, the first module comprises oxygen generation, where nitrite is reduced to nitric oxide by nitrite reductase (NIR) (Wang et al., 2019), followed by the action of nitric oxide dismutase (NOD) to yield N_2_ and O_2_. The second module encompasses the oxidation reactions, where AlkB would be used to oxidize FAs and FAOHs to ω-hydroxyacids and diols, respectively, using rubredoxin as electron carrier (Figure 3B). The remaining portion of the AlkBGT complex is responsible for regenerating reduced rubredoxin by consuming NADH. Further oxidation of ω-hydroxyacids to dicarboxylic acids, however, would require additional enzymes, like cinnamyl alcohol dehydrogenase (yjgB) and 6-oxohexanoate dehydrogenase (chnE) (Clomburg et al., 2015), even though it is reported that AlkBGT complex can further oxidize hydroxyls to carboxyls and even ester groups, a phenomenon called overoxidation (van Nuland et al., 2017). Protein discovery and engineering will play a key role in bringing these pathways to life in the chain elongating bacteria. For example, understanding anaerobic carboxylation mechanisms and discovering other related pathways would circumvent the need of using oxygen and microcompartments. Additionally, to the best of our knowledge, no FAOHs have been used as substrates for AlkBGT, which would require protein characterization and engineering to achieve diol production if intra-aerobic conditions are necessary.

The reactions described above were incorporated into our model and the effect on redox balance and ATP yield was assessed for the production of adipic acid and 1,6-hexanediol from hexanoic acid. NO_2_^−^ was supplied in our simulations as a substrate to source O_2_ for the hydroxylation step. Nitrite reduction via NIR requires an investment electron equivalents from NADH. However, the model predicted that the production of adipic acid does not require additional electrons equivalents from H_2_ nor lactate as H_2_ evolution and molar yield of adipic acid from lactate was the same as the hexanoic acid case. For adipic acid, the ω-hydroxyacid intermediate that is produced by AlkBGT is subsequently oxidized, releasing an NADH that is required for the reduction of NO_2_^−^ and balancing redox. However, in the case of 1,6-hexanediol, an additional electron source is required for the initial reduction of hexanoic acid to hexanol. Thus, diol production is not as carbon efficient because additional electron equivalents from lactate are required to reduce both terminal ends into alcohols, leading to the co-production of acetate to balance carbon, yielding a net increase in ATP yield due to substrate-level phosphorylation through PTA/ACK. ECM found that 1,6-hexanediol carries an enzyme cost marginally lower than that of hexanol produced via AOR, 2.5 times the cost of butyrate. Cost decreases since both pathways share an ATP yield of 0.5, while diol production does away with the CoAT reaction. This reaction is itself enzymatically expensive due to its low driving force, and it increases the cost of PTA and ACK reactions as CoAT requires a higher acetate concentration. Adipic acid production requires 7.5 times more enzyme investment per ATP flux than butyrate (Figure 4). This suggests that it may be more challenging to achieve high adipic acid productivity due to a reduction in growth rate. On the other hand, 1,6-hexanediol production could even increase growth rate vs. hexanoic acid, similarly to hexanol production.

#### Methyl ketone Production

Methyl ketones (MKs) are another chemical class that can be produced using MCFAs as precursors. Currently, in the engineered RBO cycle, methyl ketones are obtained from the hydrolysis of oxoacyl-CoA molecules by the thioesterase FadM. This reaction initially generates ketoacids, which in turn spontaneously decarboxylate to result in the final MK (Figure 3C) (Yan et al., 2020). In addition to FadM introduction, other metabolic engineering strategies are necessary to achieve methyl ketone production in chain elongating bacteria. The first strategy would be to substitute the native CoATs by FadM, translocating termination from acyl-CoA to oxoacyl-CoA. Since the RBO cycle is essential to redox balance and ATP production in chain elongating bacteria, sole acetone and 2-butanone production would not be possible, as FadM would terminate before one complete turn of the RBO cycle. Therefore, to optimize MK production, it is necessary to allow MCFA with MK co-production.

FadM was incorporated into our model to assess the potential to produce 2-pentanone from 6-oxohexanoyl-CoA. The model predicted that 2-pentanone production is not optimal for ATP yield which aligns with the early termination initiated by FadM at 6-oxohexanoyl-CoA, where the ATP from the complete elongation to hexanoic acid is not produced. To assess the effect of 2-pentanone production on ATP yield, we fixed 2-pentanone production at various values and determined the overall pathway stoichiometry at each point using pFBA with ATP yield set as the objective function (Table 3). It was found that 2-pentanone is negatively coupled with butyrate and ATP production (Figure 5). Further, 2-pentanone production is thermodynamically feasible only for MK production yields less than 0.15 mol 2-pentanone per mol lactate. Since 2-pentanone production is feasible in *Escherichia coli* (Lan et al., 2013), it suggests that the concentrations for standard conditions are likely incorrect to estimate the pathway driving force. CO_2_ production also increases with increasing 2-pentanone production due to the spontaneous decarboxylation of the ketoacid. In isolation, this loss of CO_2_ severely impacts the carbon efficiency of this production pathway. However, it could be alleviated if methyl ketone production was conducted in the context of a microbial community, where other functional guilds could recycle the CO_2_ (Baleeiro et al., 2023).

**Figure 5.**
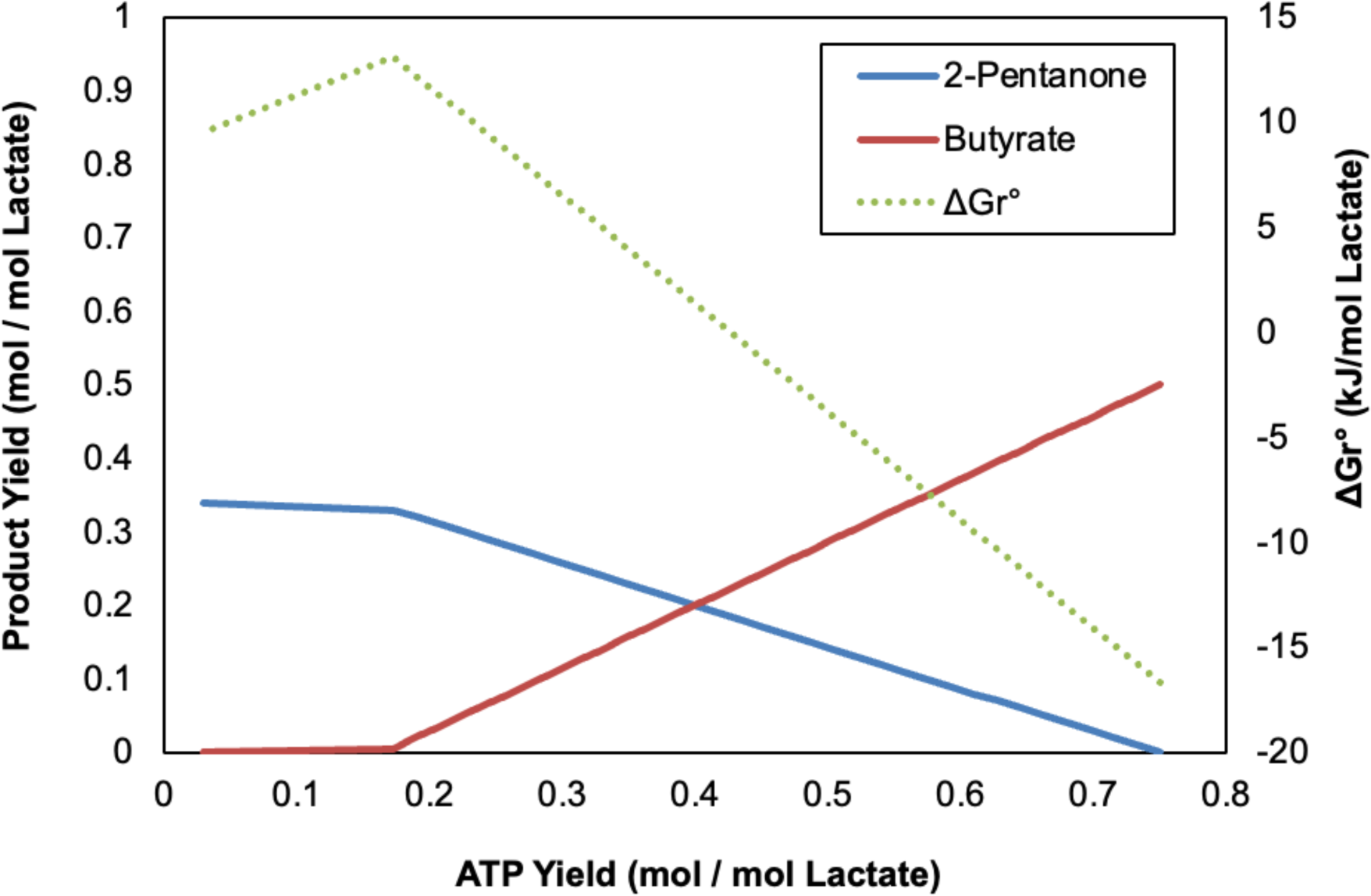
Production profiles for the co-production of 2-pentanone and butyrate from lactate and standard Gibbs free energy of reaction. Molar yields of ATP are negatively related to increasing production of 2-pentanone. Co-production of butyrate is required with increasing 2-pentanone production.

**Figure 6.**
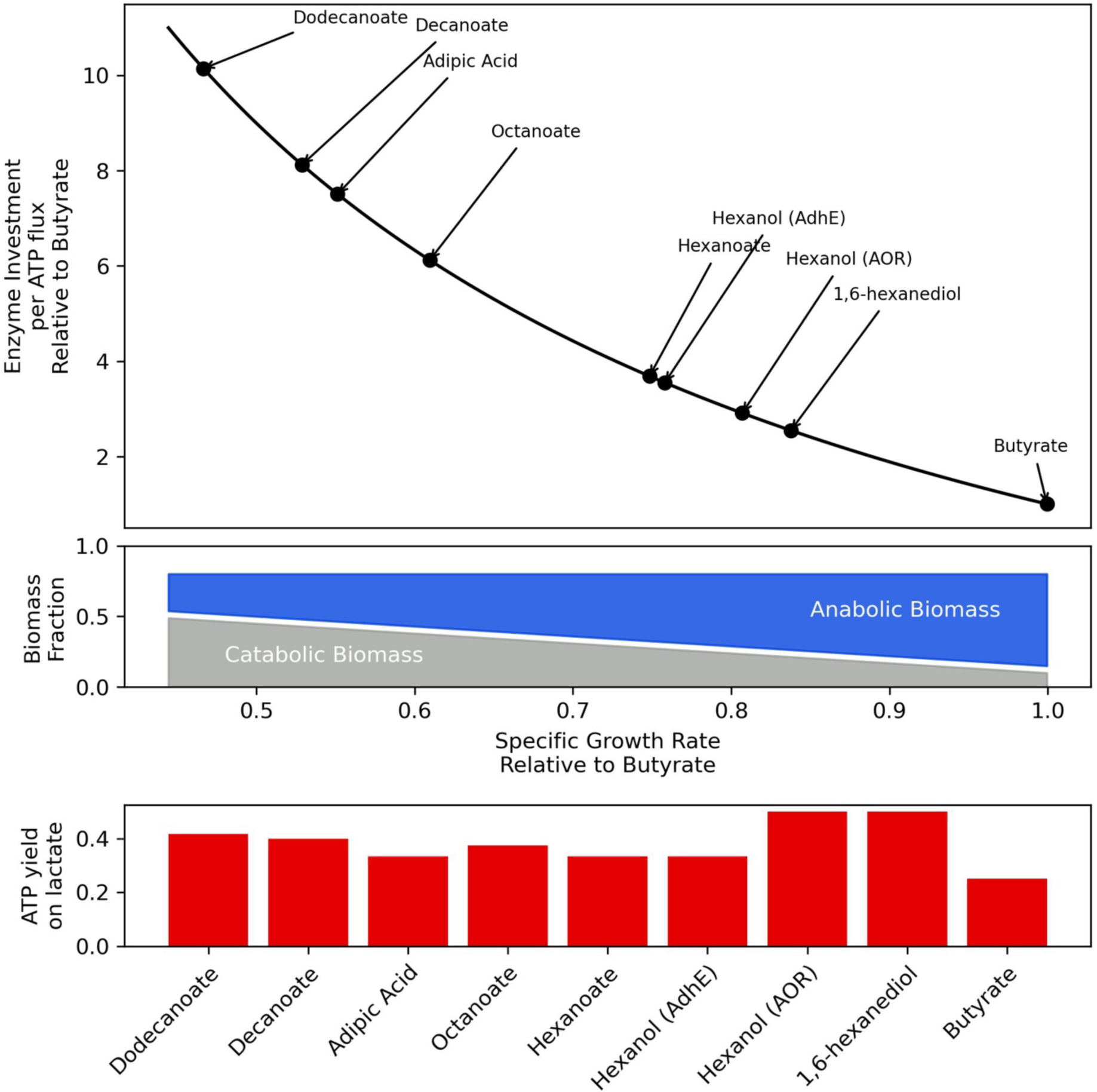
Top: Enzyme cost per ATP flux relative to butyrate vs. specific growth rate relative to butyrate, as derived from a simplified resource allocation model (Supplementary text). Middle: Anabolic fraction of biomass corresponding to specific growth rate (blue) and catabolic fraction of biomass remaining to supply ATP at a given growth rate. Bottom: ATP yield on lactate for different chain elongation simulated by pFBA.

**Table 3.**
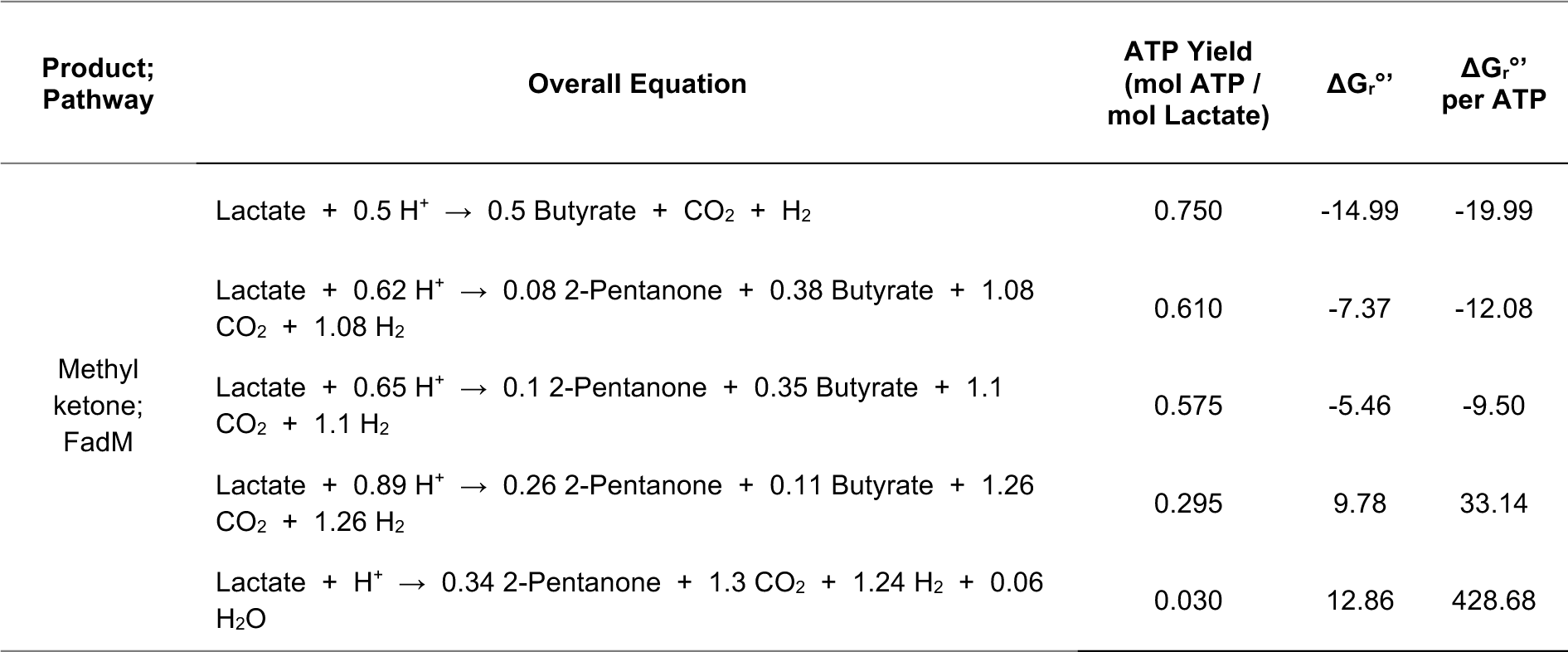
Selected predicted pathway stoichiometries for 2-pentanone production.

| Product;<br>Pathway | Overall Equation | ATP Yield<br>(mol ATP /<br>mol Lactate) | $\Delta G_r^{\circ}$ | $\Delta G_r^{\circ}$<br>per ATP |
| --- | --- | --- | --- | --- |
| Methyl<br>ketone;<br>FadM | Lactate + 0.5 H<sup>+</sup> → 0.5 Butyrate + CO<sub>2</sub> + H<sub>2</sub> | 0.750 | -14.99 | -19.99 |
|  | Lactate + 0.62 H<sup>+</sup> → 0.08 2-Pentanone + 0.38 Butyrate + 1.08 CO<sub>2</sub> + 1.08 H<sub>2</sub> | 0.610 | -7.37 | -12.08 |
|  | Lactate + 0.65 H<sup>+</sup> → 0.1 2-Pentanone + 0.35 Butyrate + 1.1 CO<sub>2</sub> + 1.1 H<sub>2</sub> | 0.575 | -5.46 | -9.50 |
|  | Lactate + 0.89 H<sup>+</sup> → 0.26 2-Pentanone + 0.11 Butyrate + 1.26 CO<sub>2</sub> + 1.26 H<sub>2</sub> | 0.295 | 9.78 | 33.14 |
|  | Lactate + H<sup>+</sup> → 0.34 2-Pentanone + 1.3 CO<sub>2</sub> + 1.24 H<sub>2</sub> + 0.06 H<sub>2</sub>O | 0.030 | 12.86 | 428.68 |

## Conclusion

Developing chain elongating bacteria as a bioproduction platform for oleochemical synthesis could play a crucial role in sustainable manufacturing from carbon-rich wastes streams. Our analyses demonstrate that the synthesis of C_9_-C_12_ fatty acids, as well as conversion to their corresponding fatty alcohols, dicarboxylic acids, diols, and methyl ketones are thermodynamically feasible and generate sufficient ATP production for growth coupling. In the case of C_9_-C_12_ fatty acids, enzyme cost analyses point towards a trade-off between growth rate and ATP yield with increasing chain length which may explain why chain elongation in nature terminates at chain lengths of C_8_.

Other aspects that need to be investigated to understand limitations in chain length include the effect of product toxicity and enzyme-substrate affinity for longer-chain fatty acids. This includes strategies chain elongators employ to protect against MCFA toxicity and evaluation of the substrate range that can be accommodated by native RBO cycle enzymes. To address these problems, genetic engineering tools need to be developed, including genetic cargo delivery systems, the construction of regulatory element libraries and gene knock-in/knock-out methods to delete native chain elongating bacteria genes and to enable heterologous gene expression. These genetic modifications will allow the production of a broad range of oleochemicals through chain elongation using the pathways proposed in this work. The production of longer chain length fatty acids and their subsequent conversion into alcohols are ideal targets for initial demonstrations of oleochemical production using engineered chain elongating bacteria as these products are not redox limited in our analyses. However, the functionalization of the aliphatic end of the fatty acids required for dicarboxylate and diol production will be a significant challenge due to the anaerobic requirements of chain elongating bacteria. While we present an avenue to accomplish this through nitrite reduction and the use of microcompartments, screening of other enzymes that can functionalize fatty acids without oxygen will be important to chain elongating bacteria metabolic engineering efforts. Additionally, the production of methyl ketones was found to be feasible only for select pathway stoichiometries and not strongly growth coupled as it requires carboxylate co-production for ATP production. Given previous demonstrations of methyl ketone production in *Escherichia coli* under microaerobic conditions (Lan et al., 2013), other conditions and production pathways should be investigated to improve yields in chain elongators.

Overall, our analyses indicate that chain elongating bacteria are promising biomanufacturing chassis for accessing several oleochemical product classes. Developing genetic tools for chain elongating bacteria will be critical for understanding the fundamental physiology of this functional guild and will be required to expand their product spectrum, in isolation as pure cultures, and in the context of self-assembled and synthetic microbiomes.

## Methods

### Predicting Pathway Stoichiometry with Parsimonious Flux Balance Analysis

Python3 and CobraPy were used for the metabolic modelling. Previously, the iFermCell215 model was created to represent a generalized chain elongation microbiome with several functional guilds capable of utilizing different electron donors for elongation up to C8 (Scarborough et al., 2020). iFermCell215 was simplified by removing some inapplicable reactions and further modified with 26 metabolites and 30 reactions required for the production of C_9_-C_12_ carboxylates to produce iFermCell193. iFermCell193 was further constrained to capture the metabolism of typical CEB, in alignment with the proposed stoichiometric models previously described to obtain a generalized core model for chain elongating bacteria (Angenent et al., 2016; Scarborough et al., n.d.). Using iFermCell193, the production of fatty acids from lactate (main text), ethanol, glucose, xylose, and glycerol (Supplementary Material) was modelled with parsimonious enzyme usage flux balance analysis (pFBA) (Lewis et al., 2010) where net ATP yield was set as the objective function, used here as a proxy for growth.

iFermCell193 was further modified with 70 metabolites and 163 reactions to produce iFermCell356 based on the proposed production pathways for expanding the chain elongating bacteria product spectrum to produce fatty alcohols, dicarboxylates, diols, methyl ketones. The production of these bioproducts from different electron donors was modelled with pFBA where net ATP yield was set as the objective function, except methyl ketone production.

Python package equilibrator-api (https://gitlab.com/equilibrator) was used to predict the Gibbs free energy of formation for metabolites at physiological conditions of T = 25 °C, pH = 7, pMg = 0, Ionic Strength = 0 M. To calculate overall Gibbs free energy change of reaction for the production of the different products from lactate, the Gibbs free energy change of formation and the overall pathway reaction stoichiometry predicted by the model were used. The models, configuration files, and final flux solutions are provided in the Supplementary Material.

### Enzyme Cost Minimization

ECM was developed by Flamholz et al. and performed here with the Python packages equilibrator-api and equilibrator-pathway (https://gitlab.com/equilibrator). A reversible Michaelis Menten kinetic expression is substituted with a flux-force efficiency term, 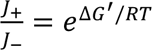, to yield an expression for the enzyme concentration [*E*] required to sustain a given flux *J* through a pathway reaction:

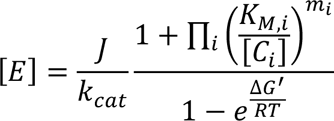

Where *k_cat_* is the enzyme’s rate constant, [*C_i_*] is the concentration of reactant i, *K_M,i_*, is the enzyme’s saturation constant for reactant i, *m_i_* is the stoichiometric coefficient of reactant i, and Δ*G*^’^ is the Gibbs free energy change, estimated via the component contribution method at pH 6 (Noor et al., 2013). This expression can be decomposed into 3 factors to facilitate interpretation: 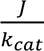 represents the stoichiometric enzyme demand; 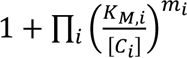 encapsulates the cost incurred when the enzyme is not saturated; 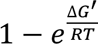 reflects the increased enzyme requirement to sustain a net flux when a reaction is near equilibrium. The total enzyme cost per unit pathway flux (gram protein/mole product/hour) is calculated by summing reaction costs multiplied by both their stoichiometry in the pathway *n_i_* and their enzyme’s molar weight *M_i_*, then dividing by the arbitrary pathway flux *J* in units of moles per hour. Total enzyme cost per unit flux is minimized by gradient descent given bounds on metabolite concentrations and the requirement for all reactions to be exergonic:

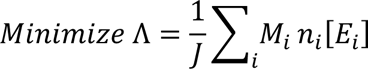

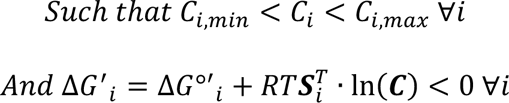

Where 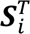 is the ith row of the stoichiometric matrix’s transpose, containing the stoichiometric coefficients for each metabolite in the ith reaction, and ln(***C***) is the vector of logarithms of each metabolite’s concentration. As a first approximation, all enzymes are assumed to share rate and saturation constants and molecular weight (k_cat_ = 200 s^−1^; K_m_ = 0.2 uM, MW = 40 kDa). Pathway specific metabolites are bounded between 1 uM and 10 mM, apart from oxoacyl-CoAs which can drop to 0.4 uM. Universal metabolites such as ATP and NADH are set to concentrations derived from literature (Bennet et al. 2009). The reduction potential of ferredoxin is set to −400 mV. Dissolved oxygen concentration was set to 6.6 uM (0.005 atm at equilibrium), which is expected to be sub-lethal to obligate anaerobes (Loesche, 1969). Dissolved hydrogen concentration was 1.4 mM (0.2 atm) and dissolved CO_2_ concentration was 10 uM. Ionic strength was set to 250 mM. A proton motive force of 200 mV was used for butyrate production, whereas 100 mV was used in all other cases, due to the reversal of RNF activity required for butyrate production without an electron acceptor.

## Acknowledgements

The authors would like to thank Dr. Adam M. Guss for their support and expertise on this project.

