## Supplementary material for "Evaluating the feasibility of medium chain oleochemical synthesis using microbial chain elongation": SI: Supplementary Figures Captions.docx

**Supplementary Figure 1:** Reaction Gibbs free energy changes in chain elongation pathways predicted by Enzyme Cost Minimization (Flamholz et al. 2013). Physiological ∆G°’ (grey) are evaluated at uniform metabolite concentrations of 1 mM. Optimized ∆G’ follow from the set of metabolite concentrations which minimizes enzyme demand per unit pathway flux. All are evaluated at pH 6 and 25 °C.

Figure 2 – Hypothetical anaerobic oxidation of heptanoic acid to adipic acid. Adapted from Heider et al., 2016. EBDH: ethylbenzene dehydrogenase; PEDH: phenylethanol dehydrogenase; APC: acetophenone carboxylase; BAL: benzoylacetate-CoA ligase; THL: thiolase. Created with BioRender.com
