## Supplementary figures and images for "Evaluating the feasibility of medium chain oleochemical synthesis using microbial chain elongation"

### Hypothetical Anaerobic carboxylation of fatty acids.png

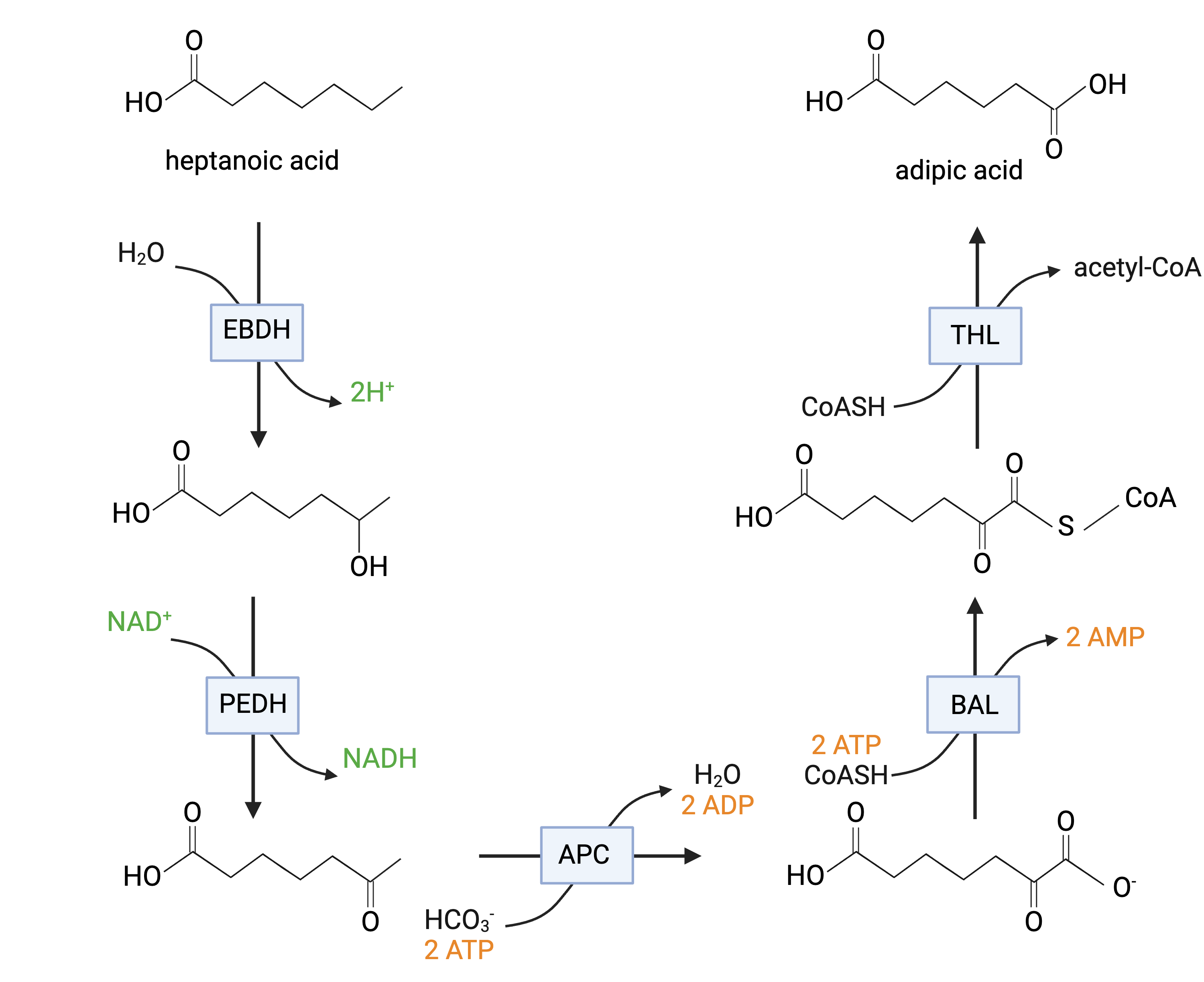

### Supplementary Figure 1.png

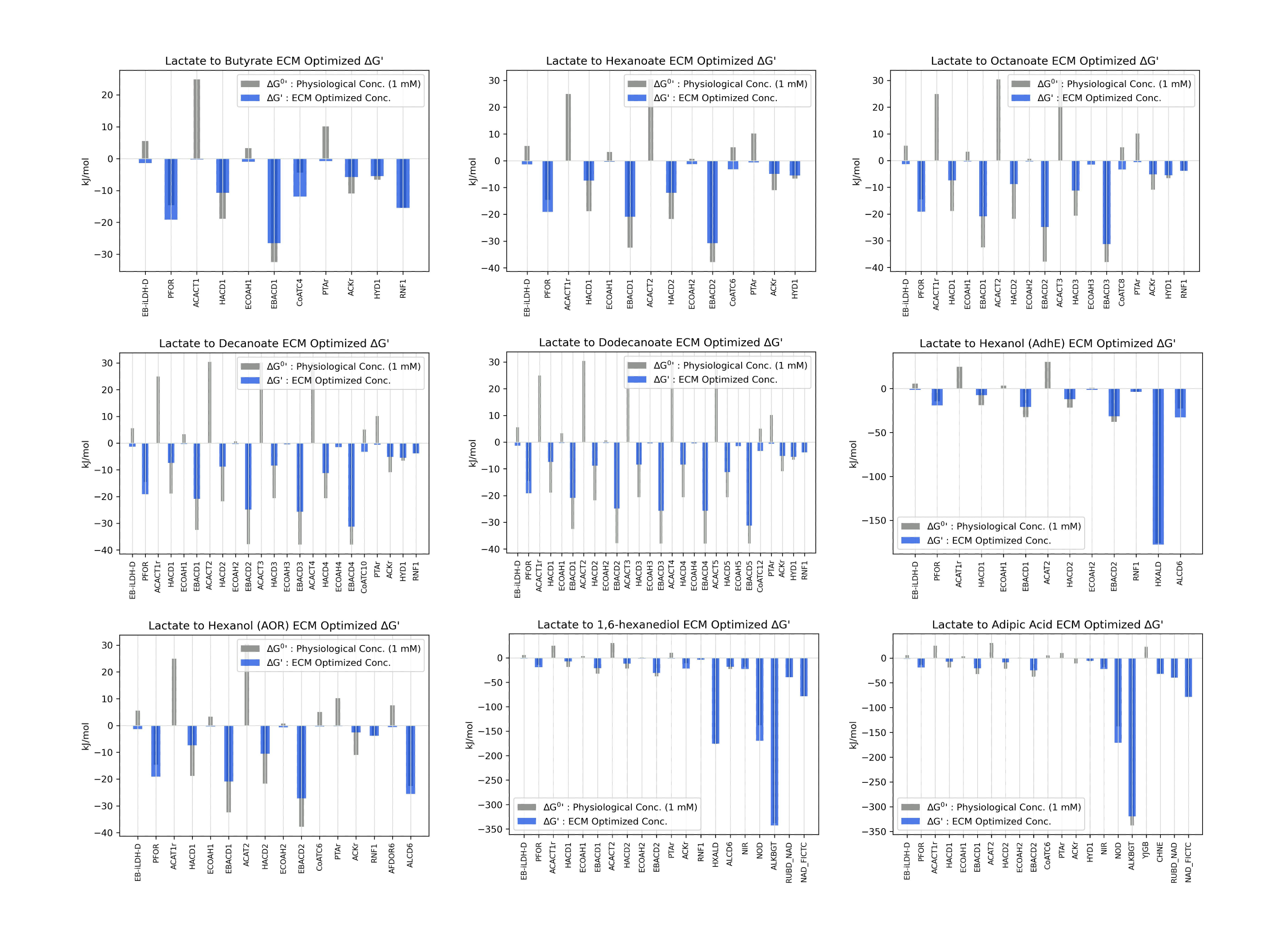
